# A Comprehensive Overview of Genomic Imprinting in Breast & Its Deregulation in Cancer

**DOI:** 10.1101/257824

**Authors:** Tine Goovaerts, Sandra Steyaert, Chari A Vandenbussche, Jeroen Galle, Olivier Thas, Wim Van Criekinge, Tim De Meyer

## Abstract

Genomic imprinting, the parent-of-origin specific monoallelic expression of genes, plays an important role in growth and development. Loss of imprinting of individual genes has been found in varying cancers, yet data-analytical challenges have impeded systematic studies so far. We developed a mixture distribution model to detect monoallelically expressed loci in a genome-wide manner without the need for genotyping data, and applied the methodology on TCGA breast tissue RNA-seq data. We identified 35 putatively imprinted genes in healthy breast. In breast cancer however, HM13 was featured by significant loss of imprinting and expression upregulation, which could be linked to DNA demethylation. Other imprinted genes (25 out of 35) demonstrated consistent expression downregulation in breast cancer, which often correlated with loss of imprinting. A breast imprinted gene network, deregulated in cancer, might hence be present. In summary, our novel methodology highlights the massive deregulation of imprinting in breast cancer.

---

Breast cancer is the most common type of cancer in women^1^. It is a very heterogeneous disease with major differences in incidence, clinical outcome, prognosis and response to therapy^2,3^. Gene expression profiling led to the division of breast cancer in five different molecular subtypes: Luminal A, Luminal B, HER2-enriched, Basal-like and Normal-like^2,4^. These subtypes differ amongst others in expression of the oestrogen receptor, progesterone receptor, human epidermal growth factor receptor 2 (HER2) and in histological grade^3^.

An early occurring aberration in cancer is loss of imprinting (LOI)^5^. Imprinting refers to the monoallelic expression of genes in a parent-of-origin specific manner. In diploid eukaryotic organisms, the maternal and paternal copies of most genes are expressed at similar levels. For imprinted genes, however, only a single allele is transcriptionally active^6–8^. Imprinting patterns may vary between tissues^9^. Imprinted genes are mostly clustered and regulated by imprinting control regions (ICR), which are typically under DNA methylation control, though also H3K27me3 was recently demonstrated to be involved^10,11^. Imprinted genes play an important role in development and placental biology^12^. Furthermore, as dosage of imprinted genes is crucial, disruption of imprinting can result in a number of human imprinting syndromes and may predispose to cancer by promoting tumourigenic or suppressing antitumour mechanisms^13–16^. Some well-known diseases are Angelman Syndrome (functional loss of the maternal, active allele of UBE3A), Prader-Willi Syndrome (loss of the paternal, active allele of SNRPN) and Beckwith-Wiedemann Syndrome (LOI on chromosome 11)^14,17^.

LOI results in biallelic expression, typically due to activation of the silent allele. Indeed, recent experiments in mice demonstrated that demethylation at imprinted genes leading to LOI made cells susceptible to cellular transformation and tumourigenesis^18^. For instance, aberrant biallelic expression of the imprinted IGF2 locus is thought to promote tumourigenesis by inhibiting apoptosis in colorectal cancer^19^ and to lead to over-proliferation defects in lung, colon and ovarian cancer^20^. LOI of other imprinted genes, such as H19, PEG3, MEST and PLAGL1, was also discovered in varying cancers^21^.

However, several studies suggest a far more complicated story, where LOI is associated with silencing of the normally active allele^5^. For example, recent studies identified major expression downregulation of reportedly imprinted genes in cancer^22,23^. Moreover, in a oesophageal cancer study, LOI of IGF2 was specifically associated with downregulation of IGF2 expression, and improved survival^24^. Also in prostate cancer, no increased expression was found for IGF2 despite LOI^25^. Notwithstanding the major relevance of LOI in cancer, this fragmentary evidence demonstrates that the current paradigm of the role of LOI in cancer (i.e. growth & tumour promoting expression) requires additional evaluation. A recent study by Ribaraska *et al.* found downregulation of several imprinted genes in prostate cancer, but stable DNA methylation^23^. These results imply the existence of an imprinted gene network in which these genes are co-regulated, as was also observed in mice^26^. However, only a small subset of the imprinted genes was analysed in the latter study.

Systematic analyses of the impact of LOI are still lacking. Indeed, although monoallelic expression is a well-investigated topic, only few regions are well-characterised in humans, and only a single study thoroughly evaluated tissue specific imprinting patterns^9^. Also, to date, most effects of aberrant monoallelic expression on cancer have been studied at single imprinted loci^18^. Moreover, the practical applicability of existing high-throughput methods is greatly hampered by the necessity for genotyping next to (typically) RNA-seq data. Thus, there is a need to systematically profile (i) monoallelically expressed/imprinted loci and (ii) their deregulation (LOI) in cancer, preferably solely based on RNA-seq data.

Here, such a methodology was developed and – given indications for massive differential expression of imprinted genes in this tumour^22^ - applied on TCGA breast case/control data, leading to the identification of 35 putatively imprinted genes in breast of which 8 were featured by LOI in at least one breast cancer subtype. Comparison with whole exome sequencing (WES) data demonstrated that (i) the RNA-seq based results were generally reliable, and (ii) that avoiding the use of WES data leads to a far higher genome-wide character. Intriguingly, the results indicate that LOI, also when linked to survival (ZDBF2), is more often associated with lower expression than higher expression of the corresponding locus, though exceptions exist (e.g. HM13). Overall, our results demonstrate that deregulation of imprinting is an important feature in breast cancer, and likely in other human cancers, but is not automatically associated with higher gene expression. Furthermore, it underlines the efficacy of the proposed strategy for the identification of imprinted regions and their deregulation.

## RESULTS

### Detection of imprinting in healthy breast tissue

First, a novel methodology was developed and applied to screen for imprinted SNPs in RNA-seq data. Contrasting previous genome-wide methods, no DNA genotyping data is required, as we solely rely on genotyping of RNA-seq data. The basic rationale is that in case of 100% imprinting, no heterozygous samples can be found in RNA-seq data, as they perfectly resemble homozygous samples (only a single allele is expressed) (see Methods section and Steyaert et al.^27^). The established imprinting model describes the data for each SNP as a mixture of homozygous and heterozygous samples, more specifically as a mixture of genotype dependent binomial distributions, with weights derived from Hardy-Weinberg equilibrium. The model has been developed to allow for sequencing errors and partial imprinting. Indeed, one parameter describes the degree of imprinting, and it can be evaluated whether this parameter is significantly higher than 0 using a likelihood ratio test.

Upon application on 113 TCGA breast control samples, 127 SNPs were considered to be possibly imprinted (false discovery rate (FDR) < 0.05), and dbSNP annotation was found for 125 SNPs. The 125 possibly imprinted SNPs corresponded with approximately 35 genes, of which at least 16 were already known to be imprinted. Note that annotation of the SNPs to specific genes was often difficult as many overlapping genes were found (see Supplementary Results, Section 1). For example, MCTS2P is a retrogene copy and located in HM13, making it uncertain in which gene the detected SNP was located. Table 1 lists the identified SNP loci and corresponding genes. All alleles of the imprinted SNPs corresponded to dbSNP listed alleles (data not shown). As examples, the resulting mixture distributions of IGF2 and SNRPN are shown in Figure 1(a) and Figure 1(b), respectively. The distributions show that these loci are clearly depleted of heterozygous samples, corresponding with possible imprinting. Moreover, Figure 1(a) indicates that IGF2 imprinting is only partial, underscoring the suitability of the flexible distribution model used here. Similar figures for the other genes can be found in Supplementary Figure 1-11.

**Figure 1.**
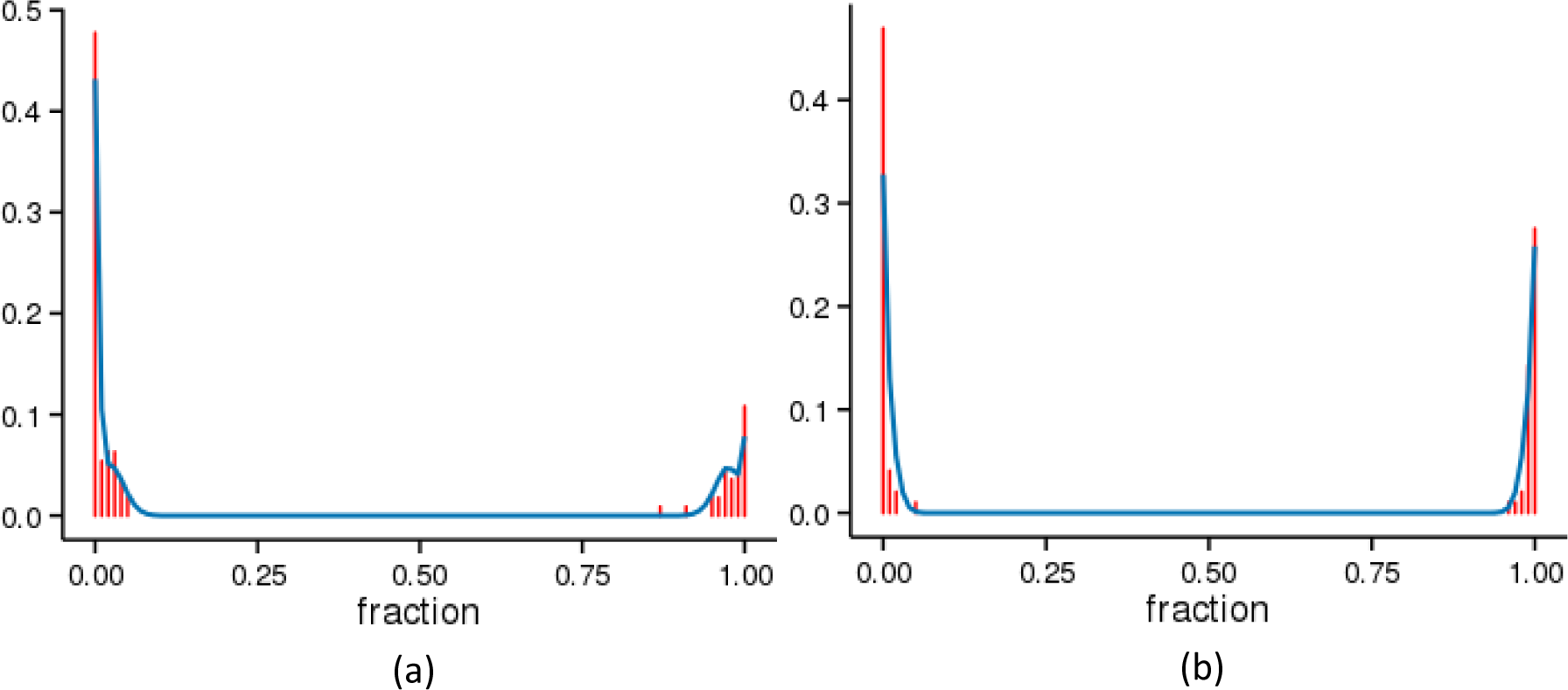
Observed (red) and modelled (blue) fraction of alternative alleles for two significantly imprinted SNP positions, i.e. (a) IGF2 (rs2585, adj. p-value < detection limit) (b) SNRPN (rs705, adj. p-value = 1.81E-71)

**Table 1.** Genes featured by monoallelic expression in breast tissue. The columns show the gene symbol (Gene symbol) and the ID of significantly imprinted SNP locus falling in these genes (SNPs) with its FDR-adjusted p-value between brackets. Genes which are known to be imprinted or for which there is prior evidence of imprinting are marked in red.

| Gene symbol | SNPs | Gene symbol | SNPs | Gene symbol | SNPs |
| --- | --- | --- | --- | --- | --- |
| MTCO1P12 | rs112232512 (1.42E-75) | MEG3 | rs35458454 (5.16E-21) | PWAR6 | rs62001982 (1.09E-19) |
| LINC01139 | rs61746209 (6.14E-29) | MEG3 | rs35431412 (1.62E-29) | PWAR6 | rs1045935 (6.76E-29) |
| ZDBF2 | rs7582864 (1.02E-21) | MEG3 | rs10147988 (1.07E-26) | SNHG14 | rs11637436 (1.44E-11) |
| ZDBF2 | rs3732084 (6.54E-26) | MEG3 | rs3087918 (5.47E-27) | SNHG14 | rs4344720 (7.48E-20) |
| ZDBF2 | rs1975597 (7.97E-30) | MEG3 | rs3087917 (1.64E-30) | SNHG14 | rs3863396 (2.68E-19) |
| ZDBF2 | rs1448902 (2.02E-17) | MEG3 | rs3742391 (2.13E-19) | SNHG14 | rs62002013 (3.83E-12) |
| ZDBF2 | rs4673350 (2.56E-23) | MEG3 | rs12897172 (2.31E-17) | SNHG14 | rs2356294 (4.96E-24) |
| PAX8-AS1 | rs7585510 (0.0068) | MEG3 | rs1884540 (1.78E-26) | SNHG14 | rs1043164 (6.41E-12) |
| PTX3 | rs73158510 (5.44E-07) | MEG3 | rs2400941 (1.64E-29) | SNHG14 | rs691 (6.11E-33) |
| CPHL1P | rs12497062 (0.00027) | MEG3 | rs77658190 (2.45E-17) | SNHG14 | rs13526 (3.74E-29) |
| ATP8A1 | rs11940243 (3.21E-08) | MEG3 | rs10132552 (4.26E-17) | PLIN1 | rs4578621 (3.29E-06) |
| NAP1L5 | rs8605 (3.77E-22) | MEG3 | rs3194464 (1.43E-23) | ZNF597 | rs37822 (1.47E-18) |
| NAP1L5 | rs710834 (9.10E-21) | MEG3 | rs11160606 (1.53E-23) | ZNF597 | rs37823 (3.08E-18) |
| ZNF300P1 | rs17800987 (4.87E-11) | MEG3 | rs1950628 (1.01E-26) | ZNF597 | rs11639510 (3.42E-25) |
| PLAGL1 | rs2328535 (3.09E-10) | MEG3 | rs1053900 (4.91E-34) | ZNF597 | rs37824 (3.21E-18) |
| PLAGL1 | rs9373409 (1.40E-27) | MEG3 | rs1054000 (7.58E-27) | ZNF597 | rs12737 (1.01E-21) |
| PLAGL1 | rs73006222 (2.41E-15) | MEG3 | rs8013873 (6.63E-27) | TPSB2 | rs77309587 (1.52E-08) |
| PLAGL1 | rs17615967 (1.76E-14) | MEG3 | rs11859 (2.17E-25) | USP32P2 | rs141915702 (0.0093) |
| PLAGL1 | rs77203559 (3.76E-15) | MEG3 | rs74080162 (1.15E-24) | MTRNR2L1 | rs3931649 (3.16E-28) |
| PLAGL1 | rs9321953 (1.48E-21) | MEG3 | rs4378559 (8.76E-25) | MTRNR2L1 | rs113014658 (6.82E-09) |
| LOC100294145 | rs241407 (2.71E-07) | MEG3 | rs12890215 (8.74E-25) | MTRNR2L1 | rs113626706 (5.74E-12) |
| PEG10 | rs35237090 (1.69E-17) | MEG3 | rs55996894 (5.63E-21) | MTRNR2L1 | rs3931650 (7.67E-29) |
| PEG10 | rs13073 (3.35E-13) | MEG3 | rs3742390 (6.24E-31) | ZNF331 | rs113983639 (1.99E-13) |
| PEG10 | rs7810469 (6.18E-29) | MEG3 | rs4906022 (8.42E-204) | ZNF331 | rs8110350 (4.12E-106) |
| MEST | rs10863 (<detection limit) | SNRPN | rs2554426 (9.39E-16) | ZNF331 | rs8110538 (1.74E-109) |
| HOTAIRM1 | rs706018 (5.67E-36) | SNRPN | rs705 (1.81E-71) | ZNF331 | rs8109631 (8.52E-175) |
| H19 | rs2075745 (6.63E-34) | SNHG14 | rs2732028 (3.33E-15) | PEG3 | rs4801386 (0.00011) |
| H19 | rs2075744 (3.21E-27) | SNHG14 | rs74335291 (5.12E-24) | PEG3 | rs1558355 (0.00027) |
| H19 | rs2839698 (3.18E-57) | SNHG14 | rs2732029 (6.71E-28) | PEG3 | rs723082 (1.62E-31) |
| H19 | rs2067051 (1.94E-57) | SNHG14 | rs765438 (1.96E-28) | PEG3 | rs3143 (7.86E-06) |
| H19 | rs2839701 (3.29E-61) | SNHG14 | rs2732030 (9.43E-12) | PEG3 | rs1055359 (9.07E-20) |
| H19 | rs2839704 (1.40E-44) | SNHG14 | rs10451029 (6.14E-22) | PEG3 | rs11666110 (8.06E-28) |
| H19 | rs2839702 (2.94E-50) | SNHG14 | rs2052723 (1.14E-25) | PEG3 | rs1860565 (1.02E-21) |
| H19 | rs2839703 (1.10E-46) | SNHG14 | rs2554419 (7.96E-31) | PEG3 | rs33931963 (6.73E-28) |
| H19 | rs10840159 (2.28E-50) | SNHG14 | rs719704 (3.32E-22) | HM13 | rs6058058 (1.33E-10) |
| H19 | rs3741219 (1.01E-107) | SNHG14 | rs34316840 (1.13E-25) | HM13 | rs6059869 (4.04E-14) |
| RP11-109L13.1 | rs201284359 (4.97E-10) | SNHG14 | rs2732031 (1.23E-26) | HM13 | rs6059873 (1.29E-19) |
| IGF2 | rs7873 (1.08E-268) | PWAR6 | rs2732041 (1.60E-12) | HM13 | rs6059874 (4.88E-14) |
| IGF2 | rs2585 (<detection limit) | PWAR6 | rs2732043 (1.78E-13) | HM13 (MCTS2P) | rs1115713 (7.94E-09) |
| GLIPR1/KRR1 | rs1056905 (1.20E-06) | PWAR6 | rs2732044 (1.42E-26) | GNAS/GNAS-AS1 | rs1800900 (5.02E-16) |
| DLK1 | rs1802710 (2.68E-28) | PWAR6 | rs1030389 (1.68E-28) | BCR | rs550197 (7.93E-50) |
| MEG3 | rs78793760 (4.16E-21) | PWAR6 | rs62001981 (1.41E-26) |  |  |

Next to imprinting, the model detects random monoallelic expression, as this can only be discriminated from imprinting by means of family data, unavailable in TCGA. As a consequence, several HLA, HLA-DR and HLA-DQ genes - important players in immune reactions known to be regulated by random monoallelic expression^9^ - were also identified by our methodology. Therefore, these genes were excluded from further analyses. The remaining genes (Table 1) represent the final list of putatively imprinted loci in normal human breast tissue further analysed throughout this manuscript.

As typically only few SNPs per gene were found to be indicative of imprinting, we also evaluated the other SNPs in identified genes. Often SNPs were located in intronic regions or 5’UTRs where the coverage is too low for accurate detection of imprinting. Other undetected SNPs in exons or 3’UTRs were typically featured by a high sequencing error rate, low minor allele frequency or inferior goodness-of-fit to the model. As a result, most of these “missed” SNPs were filtered out prior to application of the imprinting likelihood-ratio-test, or exhibited non-significant results due to aforementioned problems. In Supplementary Table 14 all SNPs in HM13 are shown, which clearly demonstrates that indeed mostly intronic SNPs with low coverages are missed.

### Validation of putatively imprinted regions

To independently validate our methodology, we compared these loci with the results found by Baran *et al.^9^.* By using RNA-seq data combined with genotype data from 27 primary breast tissue samples (Genotype Tissue Expression (GTEx) project), they were able to establish (partial) imprinting in breast tissue in 15 genes, including both novel and previously identified genes (Supplementary Data5, see^9^). They also detected several HLA genes, but as already stated in the previous paragraph, these are not taken into account. In summary, 13 genes were also found to be imprinted in breast tissue by our methodology, whereas 1 (= PPIEL) was not detected due to too low coverage in our samples (i.e. was filtered out prior to statistical analysis). For a second gene, SNURF, intronic SNPs were detected to be imprinted in TCGA as well. Yet, based on manual curation, it was deemed more likely that SNRPN was the imprinted gene rather than SNURF. Of the 23 genes not identified by Baran *et al.* as imprinted, 5 have been identified as imprinted in blood by Joshi *et al*.^28^. More genes concurred between the latter method and ours, however, many were curated due to annotation difficulties. Also, several genes (e.g. PWAR6 and SNHG14) were located near known imprinted regions, and for several others fragmentary evidence is available from literature.

Subsequently, TCGA whole exome sequencing (WES) data were used to evaluate whether heterozygous samples for putatively imprinted loci were indeed featured by monoallelic expression. Though low WES coverage for many (putatively) imprinted SNPs largely complicated validation - supporting the value of the genotyping-free approach introduced here - imprinting could be verified where technically feasible (see Supplementary Figure 13).

### Loss of imprinting in breast cancer

To examine possible deregulation of imprinting in cancer, alterations in allelic expression patterns of imprinted genes were investigated in breast cancer. LOI in cancer is defined here as a higher degree of biallelic expression compared to control data. We determine biallelic expression per sample by the allelic ratio (allele count with lowest expression/allele count with highest expression), which varies from 0 (perfect monoallelic expression) to 1 (perfect biallelic expression) and is independent of expression level differences in cancer. As samples with a high allelic ratio represent heterozygotes, looking at the 2P_A_P_T_ highest fraction of samples (cf. Hardy-Weinberg theorem) allows us to only take the most likely heterozygotes into account. LOI in cancer for a specific SNP was thus defined as a significant difference of the allelic ratio’s between putative heterozygous cancer and control samples (see Methods, Section 5). Though only possible for a limited set of the data, this allelic ratio was verified to be a good measure for LOI using WES data (Supplementary Figure 14).

When considering the full set of 506 breast cancer samples, three SNP loci with significant LOI, i.e. re-expression of the silenced allele, could be identified (FDR < 0.1; Table 2). These SNPs correspond to three genes, namely MEST, H19 and HM13. Figure 2 shows the mixture distributions of these genes for both the 113 control and 506 tumour samples. The plots demonstrate that for MEST (and to a minor extent for HM13) also some control samples are featured by LOI, indicating that this locus is less stringently imprinted and clearly lost its imprinting signature in breast cancer.

**Figure 2.**
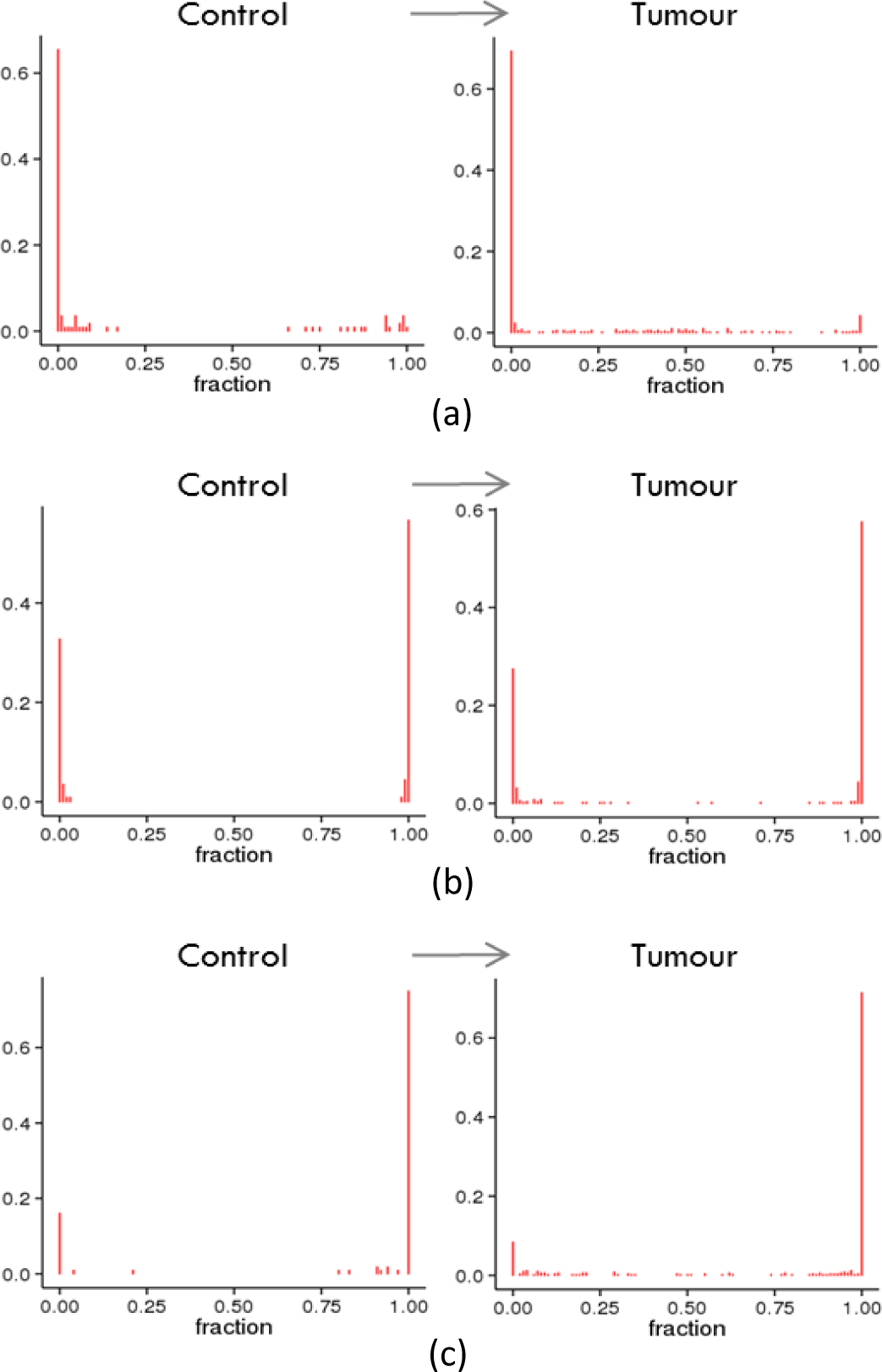
SNP positions differentially imprinted between normal and cancer samples. (a) *MEST* (rs10863, adj. p-value = 0.013). (b) *H19* (rs2839704, adj. p-value = 0.073). (c) *HM13* (rs6059873, adj. p-value = 0.067).

**Table 2.**
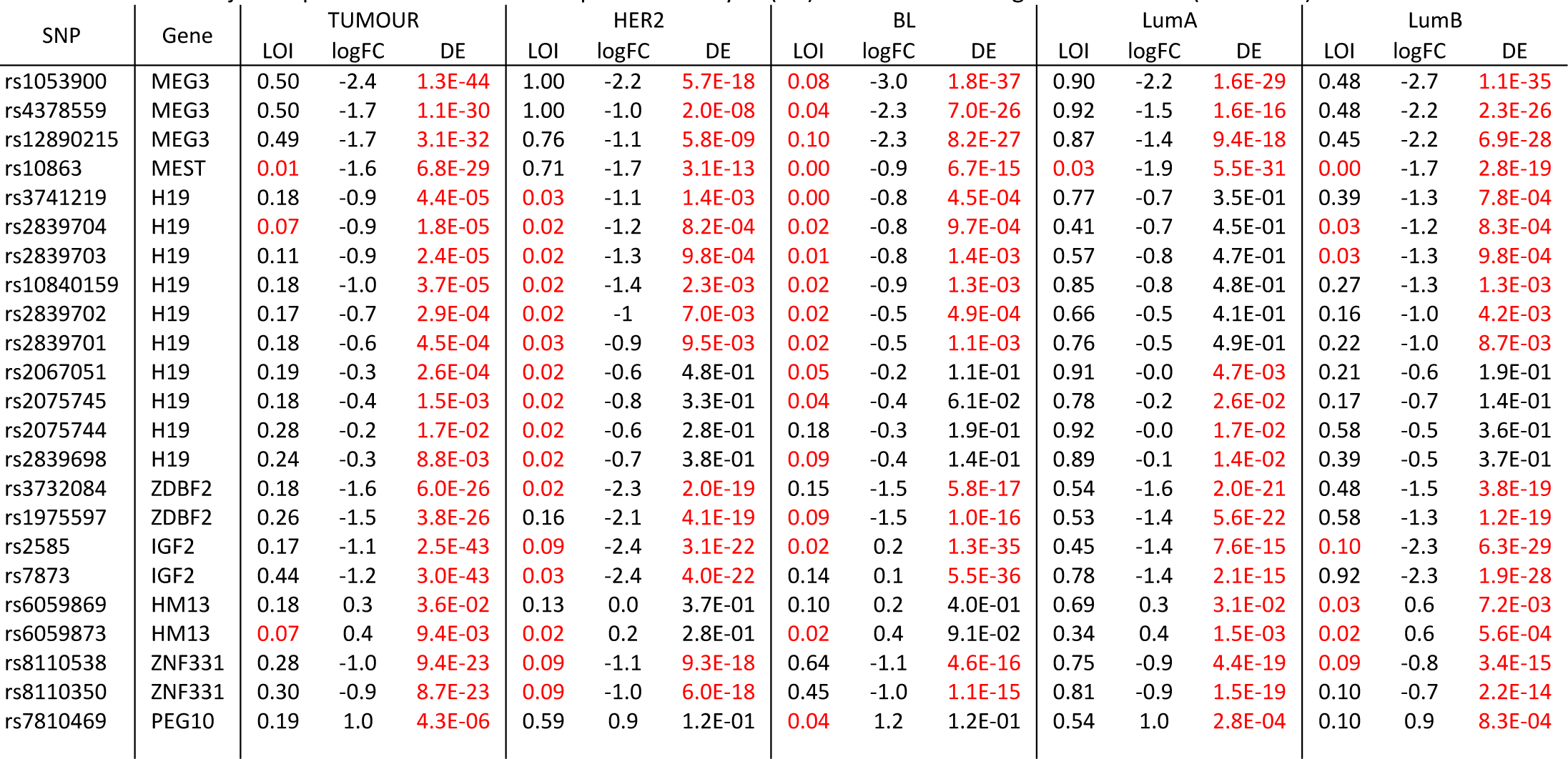
SNPs with significant loss of imprinting in control samples versus breast cancer and the different subtypes. FDR adjusted p-values are shown in column LOI with significant results (FDR<0.1) coloured in red. Also log fold changes (logFC) and FDR adjusted p-values of differential expression analysis (DE) are shown with significant results (FDR<0.05) in red.

Subsequently, the different breast cancer subtypes, namely Basal-like (BL), HER2-enriched (HER2), Luminal A (LumA) and Luminal B (LumB), were analysed individually for LOI compared to normal samples (Table 2, significant results are shown in red). Note that as there were only 8 Normal-like (NL) samples, this subtype was not taken into account. Compared to the results for the full set of tumour samples, significant LOI of MEST was detected at the same SNP position in all subtypes except for BL, whereas HM13 and H19 were significantly deregulated in all but LumA. As already stated higher, the MEST locus appears to be a flexible region in which also control samples show signs of LOI. In summary, most deregulation was found in HER2 and BL. The deregulated loci corresponded to 8 genes, i.e. ZDBF2, PEG10, MEST, H19, IGF2, MEG3, ZNF331 and HM13. For LumB, LOI was found in MEST, H19, HM13, IGF2, and ZNF331, whereas for LumA only one differentially imprinted locus could be identified (Figure 3). Distributions of the other loci featured by LOI in cancer subtypes are displayed in Supplementary Figure 15-18. Particularly the H19/IGF2 locus showed strong LOI in HER2 and BL samples: all 12 SNPs were found to be deregulated in BL, whereas for HER2 10 SNPs were differentially imprinted. Interestingly, HER2 subtype data also suggest MEG3 and ZNF331 LOI (3 significant SNP positions for MEG3 and 2 borderline significant positions for ZNF331).

**Figure 3.**
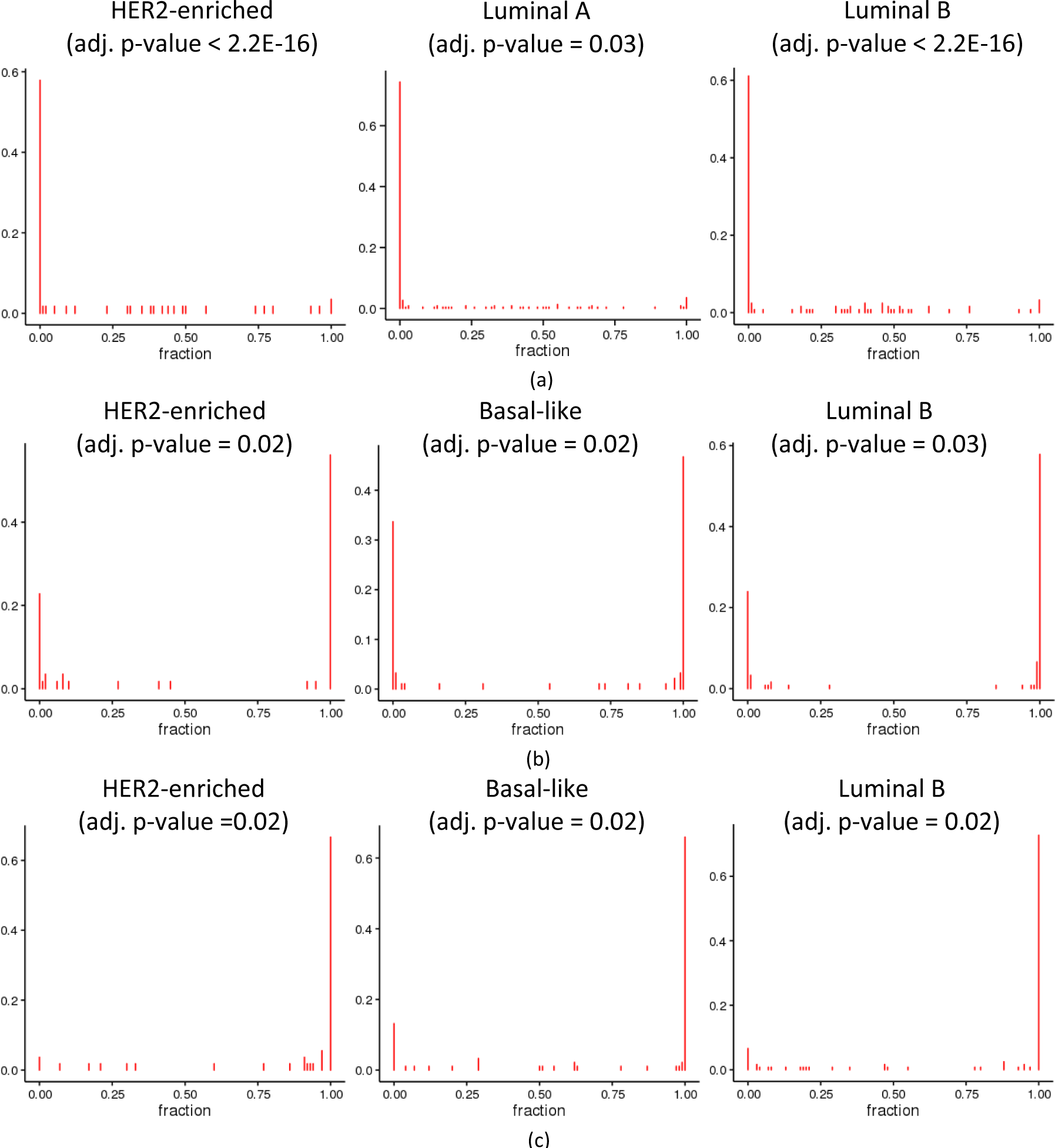
SNP positions differentially imprinted between normal and cancer subtypes, (a) MEST (rsl0863). (b) H19 (rs2839704). (c) HM13 (rs6059873).

Though most often the case, SNPs in the same imprinted gene did not always show consistent (in)significant results (Supplementary Table 15). This can typically be attributed to technical/power associated causes. For example, the SNPs displaying LOI frequently had a higher coverage than non-LOI SNPs in the same gene. Again, verification of LOI by comparison of RNA and DNA based genotypes was successful (i.e. re-expressed allele agreed with DNA based genotype), though complicated due to the low coverage WES data (Supplementary Figure 14).

Finally, it was evaluated whether LOI was associated with survival. Based on a Cox proportional hazard model (see Methods section), adjusting for age, and considering allelic ratio (allele count least expressed allele/allele count most expressed allele) as measure for LOI, ZDBF2 (2 SNPs) LOI was significantly associated with poorer survival (Figure 4, FDR-adjusted p = 0.027 and 0.086 for rs3732084 and rs1975597, respectively). If LOI was implemented categorically (dummy variable based on the allelic ratio), survival was also significantly lower with LOI of ZDBF2. Also when survival analysis was performed solely on the putative heterozygous samples (see Methods section), ZDBF2 LOI was associated with significant lower survival. The degree of LOI was not correlated to age for tumour or control data (Supplementary Table 4).

**Figure 4.**
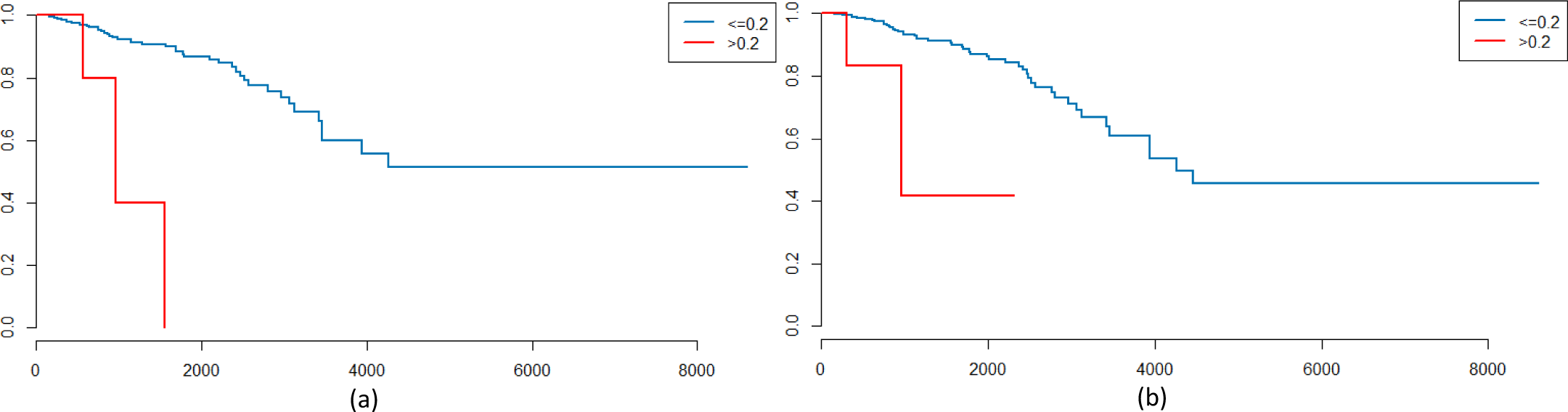
Kaplan-Meier plot of the Cox proportional hazards model for survival in function of LOI (here defined as an allelic ratio > 0.2) and age. (a) rs3732084 (ZDBF2) (b) rs1975597 (ZDBF2)

### Differential expression of imprinted genes

Imprinted genes are often differentially expressed in cancer, particularly in breast cancer^22^. in 2015, Kim *et al.* found 21 out of their 23 (91%) analysed putatively imprinted genes to be differentially expressed in breast cancer^22^. As they had compiled imprinted genes from literature irrespective of tissue type, we performed differential expression analysis in control vs tumour and control vs breast cancer subtypes for the here detected imprinted SNPs and genes. Significant differential expression (DE) was found in virtually all (99%) of the imprinted SNPs for all tumour samples. Imprinting is thus indeed heavily deregulated in breast cancer (Supplementary Table 7 and 5). Far more loci were downregulated than upregulated: 87% were detected with a negative log fold change. Again, these results correspond with the detected deregulation by Kim *et al.^22^*, but are even more clear-cut due to the prior breast specific identification of imprinted genes. The FDR-adjusted p-values and log fold changes of all SNPs showing LOI can be found in Table 2 (Supplementary Table 6 and 16 show results for all genes and SNPs, respectively).

We would expect that loss of imprinting, i.e. re-expression of the silenced allele, implies upregulation of the imprinted gene. However, at least in breast cancer, LOI is not associated with overexpression of the corresponding gene. The only clear exception was HM13, for which LOI implied higher expression in most subtypes (and IGF2 and PEG10 to a lesser extent in BL, Table 2). For the other SNPs, LOI corresponded to downregulation of expression.

We additionally verified these results in the heterozygous samples, as LOI cannot be observed in homozygotes (Supplementary Results, Section 4). Only for 6 SNPs (located in ZNF331, HM13, KRR1/GLIPR1, USP32P2, ZDBF2 and H19) sufficient WES data were available to accurately verify the LOI and DE results. In ZNF331, significant downregulation was found between the LOI tumour samples and non-LOI control samples, but also when compared to non-LOI tumour samples. HM13, on the other hand, was significantly upregulated in LOI samples (compared to non-LOI tumour as well as non-LOI control data), further confirming the results presented above. KRR1/GLIPR1 was significantly lower expressed in LOI tumour data compared to non-LOI control samples, while for the other SNPs no differences were detected. Note that the latter can particularly be explained by low power, as only few samples were featured by sufficient WES coverage for the SNPs of interest.

### LOI occurs predominantly due to silencing of the active allele

Previous research has already demonstrated that the basic concept of LOI in which re-expression of the imprinted allele leads to higher expression of imprinted genes is not always correct. Here, we found that LOI is particularly associated with downregulation of expression. We hence evaluated the association between LOI and DE into more detail.

We hypothesised that LOI may be particularly caused by the presence of partial imprinting, i.e. incomplete silencing of the imprinted allele, cf. IGF2 Figure 1(a). If expression of the normally active allele is subsequently downregulated, expression levels for both alleles will become more similar, which can be perceived as LOI. To evaluate this hypothesis, the expression of the normally silenced allele (i.e. the allele with least expression) was used as a measure for LOI. Results indicated no significant DE between cancer and controls of the “silenced” allele, supporting the hypothesis that LOI in breast cancer is particularly a by-product of the silencing of the active allele. Nevertheless, also the low number of LOI samples and thus decreased power may have an impact. It should be noted that for HM13 - clearest example of an upregulated imprinted gene - a significant (unadjusted) p-value was observed for SNP rs6059873 (higher expression of silenced allele) whereas this was not the case for the other imprinted loci (Supplementary Table 10).

### LOI, DE and differential methylation of HM13/MTCS2P locus

HM13 is the only gene in which both LOI and higher expression in cancer was identified (Table 2). However, not all SNPs in this gene showed consistent LOI and DE results. 80 SNPs were initially analysed in the full HM13 gene, yet only 5 SNPs were maintained upon initial data filtering (based on coverage, allele frequency a.o., see Supplementary Methods, Section 2.b-c and Supplementary Table 14). All of these were detected to be imprinted, 4 located in exon 3 of HM13 transcript 4, and a 5^th^ one intronic in HM13 but exonic in MTCS2P retrogene. Of the 4 detected imprinted SNPs in exon 3 of transcript 4, 3 demonstrated DE and 2 also LOI (particularly in Luminal B samples), whereas the 5^th^ SNP (exonic in MTCS2P) did exhibit LOI but not DE. Subsequently, it was evaluated whether there was evidence for transcript specific DE, as the other exons lack the informative SNPs required to evaluate LOI directly. Exon-specific expression data were downloaded and normalised with EdgeR. All exons were significantly upregulated in breast cancer as well as in the subtypes compared to control data, suggesting therefore no transcript specific effects (Supplementary Table 11).

Subsequently, ten Infinium HumanMethylation450k probes demonstrating approximately 50% methylation (see Joshi *et al.* Table S2 and MEXPRESS^28,29^) were analysed in TCGA. Methylation was significantly lower for two probes (cg18471488 - located near HM13 promoter region - and cg24607140 - located near MCTS2P and already associated with imprinting control of the latter gene^28^) in tumour compared to control samples (Supplementary Table 12). Also in the subtypes, methylation of cg18471488 was significantly lower. HER2 and LumB did not show any other differential methylation, while in BL methylation levels were lower for almost all probes (Supplementary Table 12). Only in LumA significantly higher methylation was found, which concurs with results mentioned higher that DE but no LOI was found in LumA. Methylation of probe cg18471488 was significantly correlated with expression of the last exon of transcript 4 (exon 3, possibly the UTR of this transcript) but also the full HM13 gene in the whole dataset (control and tumour data, Supplementary Table 13). In summary, these results show that LOI and DE of the HM13/MTCS2P locus is linked to DNA methylation aberrations, but that a more precise description is hampered by the resolution of the data at hand.

## DISCUSSION

Monoallelic expression of genes determined by the parental origin, called genomic imprinting, is an important feature for normal development and growth. The genome-wide evaluation of imprinting deregulation in cancer and diseases is currently hampered by a lack of appropriate data-analytical strategies. We hence developed a new methodology for the genome-wide detection of imprinting and its deregulation. After application on breast tissue, we were able to detect many imprinted genes and confirm major deregulation of imprinted genes in breast cancer and the varying breast cancer subtypes. Imprinted genes exhibited clear differential expression, particularly downregulation, in tumour samples. Strikingly, in HER2 and BL tumours, downregulation was associated with massive induction of LOI, though most likely only as a result of a higher relative expression of the (not completely silenced) imprinted allele due to silencing of the expressed allele. The major exception was HM13, exhibiting overexpression in cancer, particularly in LumB tumours, most likely due to LOI and re-expression of the normally silenced allele.

We analysed RNA-seq data of 113 breast tissue samples and 127 putatively imprinted SNPs in approximately 35 genes (after elimination of known random monoallelically expressed genes) were identified and used for further analysis. For 2 SNPs no dbSNP annotation could be retrieved, though later manual curation suggested rs2269621 to be located in the known imprinted gene L3MBTL1 (Supplementary Figure 11b). Although our methodology cannot assess whether the expression status of each allele is indeed determined by the parent of origin (which would require unavailable trio data), visual evaluation of the different mixture distribution plots as well as the clearly significant adjusted p-values strongly support at least monoallelic expression. Also, note that all genes demonstrating relevance in breast cancer had been associated with imprinting before. Finally, the observation that newly identified putatively imprinted genes demonstrate similar differential expression patterns as known ones provides more evidence for their imprinting status. Compared to the study by Baran *et al.*, which only used 27 breast samples, 13 genes were detected by both Baran’s and our method^9^. One gene was not evaluated by our methodology due to low coverage, whereas imprinted SNPs for the second gene (SNURF) were detected yet annotated as the overlapping gene (SNRPN). Of the 23 genes detected by our method only, at least 7 are known imprinted genes, also detected by Joshi *et al*.^28^, or genes located in the neighbourhood of imprinted genes. It should be noted that accuracy of these results largely depends on the accuracy of the underlying annotation. Nevertheless, inconsistent results between SNPs in the same gene were evaluated, and appeared to be mainly caused by technical reasons, i.e. low coverage of intronic SNPs, low goodness-of-fit to the model or low allele frequency, as demonstrated for HM13 in Supplementary Table 14.

Subsequently, we analysed 506 breast cancer samples for loss of imprinting. One gene, MEST, showed significant LOI and two genes, H19 and HM13, were borderline significant (FDR < 0.1). MEST is already known to show LOI in varying cancers, including breast cancer and we were here able to confirm this deregulation^21,30–32^. Aberrant H19 imprinting is often seen in cancer as well and is thought to have an important role in cancer development^33,34,21^. When taking into account the different tumour subtypes, we detected 23 SNPs (in 8 genes: ZDBF2, PEG10, MEST, H19, IGF2, MEG3, ZNF331 and HM13) exhibiting LOI in at least one subtype compared to the normal tissue samples. LOI was particularly present in BL and HER2 tumours. Many of these genes had been linked with LOI and cancer development before^21,30,35–38^. LOI was typically not associated with survival, except for 2 SNPs in ZDBF2, a zinc finger containing protein, yet additional research is required to independently validate this finding.

We found most of the imprinted genes to be downregulated in tumour samples. As canonical LOI would lead to higher expression, we suggest that LOI was observed here simply due to (constant) residual expression of the imprinted allele and downregulation of the active allele in cancer. The major downregulation of the set of imprinted genes could mean that an imprinted gene network (IGN) - as already detected in prostate cancer^23^ - is also present in breast tissue. Studies in murine and human prostate tissue detected a transcriptional network of co-regulated imprinted genes^26^, of which many were here deregulated in breast cancer as well. Moreover, PLAGL1 and H19 have been proposed as key players of the prostate IGN ^26,39–41^, and were also downregulated in breast cancer in this study. Independent of the presence of such a network, our results suggest that the massive LOI observed is the consequence of the downregulation of imprinted genes, rather than a primary event in breast cancer. Furthermore, the observation that newly identified putatively imprinted genes demonstrate similar behaviour as known ones provides more evidence for their imprinting status.

The major exception is HM13, which exhibited canonical LOI, i.e. re-expression of the normally silenced allele leading to overexpression in cancer, particularly in the LumB subtype. Both significant SNPs appear to be present in the third (and last) exon of transcript variant 4, possibly the UTR of this transcript. Though other evidence for imprinting of HM13 exists, only little information is available on its function - a signal peptide peptidase involved in the immune system^42,43^. Importantly, deregulation of HM13 - located on 20q - has been revealed in colorectal carcinoma: the often observed 20q gain in this tumour is associated with higher HM13 expression, which was demonstrated to lead to accelerated growth of the tumour^44^. Interestingly, in an imprinting study in normal blood samples by our group, HM13 also appeared to be featured by LOI and higher expression in a subset of samples, particularly in older individuals^45^. With respect to the mechanism of HM13 deregulation in breast cancer, we demonstrated that aberrant methylation in the neighbourhood of the HM13 promoter was linked to deregulation of its expression. Differential methylation may lead to different polyadenylation (as described in mice^46^) and hence varying transcripts, yet we found differential expression of all exons in HM13. Additionally, as in colorectal cancer, also 20q gain may lead to HM13 overexpression in breast cancer. It may lead to LOI as well, but note that amplification of a silenced allele does not necessarily imply activation of the latter. Further research is therefore necessary to unravel the exact mechanism(s) and consequences of (de)regulation of HM13 is breast cancer.

Throughout the manuscript, results were verified by comparison with available WES based genotyping data. However, low or absent coverage of WES for the corresponding loci led to a massive loss of imprinted SNPs that could be evaluated. This further underscores the benefits and more general applicability of the introduced methodology, which solely focuses on RNA-seq data. Also, this may explain why novel imprinted loci were found in our analysis (though often indirect evidence - e.g. imprinting already described in another tissues - was present), compared to methods where genotyping data is used to identify heterozygous samples prior to detecting imbalanced allelic expression in the latter, e.g. Baran *et al.^9^.*

Some methodological improvements could however further increase sensitivity and specificity. For example, the current method relies on Hardy-Weinberg equilibrium, but could be modified to also take into account population substructure. Additionally, the current mixture of binomial distributions could be updated to a mixture of beta-binomial distributions, as the latter captures more natural variation in expression between alleles. Nevertheless, the current study (cf. Figure 1 and Supplementary Figures), but also previous results^27^, clearly demonstrate that the proposed methodology is sufficiently robust. Another putative limitation of this study is the fact that tumour impurity, e.g. by infiltrating lymphocytes, may lead to the erroneous conclusion of LOI. Though it is impossible to fully exclude this possibility, it should be noted that many of the genes featured by LOI are also imprinted in blood cells (e.g. HM13). More importantly, both alleles are often expressed to equal extents under LOI (cf. Figure 3, samples with fractions roughly equal to 0.5), which would only be possible in case of 0% remaining tumour.

In conclusion, this study demonstrates that imprinting is indeed heavily deregulated in breast cancer, though the mechanism of its deregulation is complex. Many imprinted genes, also when linked to LOI, are downregulated in cancer. We therefore hypothesize an imprinted gene network to exist in breast tissue as well. One clear exception was found, HM13, with LOI and upregulation in cancer samples. We were able to detect these putatively imprinted genes and their deregulation with a newly developed method solely based on RNA-seq data. The effectiveness of our novel methodology and the advantage of solely using RNA-seq data, was hence also confirmed.

## METHODS

### 1. Data

RNA-seq data of 113 human healthy control and 506 diseased samples of the TCGA breast invasive carcinoma dataset were used in this study. RNA-seq BAM-files were downloaded from the prior TCGA data portal (https://portal.gdc.cancer.gov/legacy-archive/search/f). Invasive ductal carcinoma, which starts in the milk ducts of the breast, and invasive lobular carcinoma, which originates in the lobules, were both studied^47^. For all cancer samples, additional expression subtypes based on the PAM50 classifier were obtained from the UCSC cancer genome browser (8 NL, 92 BL, 228 LumA, 121 LumB and 57 HER2)^48^. In all of these samples, variants were called using Samtools mpileup/bcftools (v0.1.19)^49^.

### 2. Significance threshold

Throughout the manuscript, the Benjamini-Hochberg procedure was used for false discovery rate (FDR) estimation^50^. For detection of imprinting and differential expression analysis, an FDR of 5% was used as significance threshold. For analyses where loss of power is anticipated due to non-informative homozygous samples (e.g. loss of imprinting detection), an FDR of 10% was used.

### 3. Genotype calling

Genotype probabilities and corresponding nucleotide-read/sequencing error rates were calculated using SeqEM (v1.0)^51^ (option without Hardy-Weinberg equilibrium), a fast Bayesian genotype-calling algorithm based on the Expectation Maximisation (EM) algorithm to estimate the prior allele frequencies and the nucleotide-read error rate in an iterative way (for RNA-seq and WES data based genotyping). Note that imprinting biases RNA-seq based genotyping (i.e. less heterozygous samples will be detected), yet that allele frequency estimates are unbiased as both alleles have an equal chance to be imprinted.

### 4. Detection of imprinting

The rationale behind the proposed methodology is that biallelic expression yields RNA-seq data (or other similar sequencing data) that is in Hardy-Weinberg equilibrium (HWE) for each locus, i.e. if SNPs are present for a locus, both homozygous and heterozygous subjects will be detected at a predictable rate (under HWE assumptions)^52^. However, in case of monoallelic expression, heterozygous samples will no longer be detected in RNA-seq data resulting in deviation from the HWE, which can be measured (Figure 5(a)). Throughout the Methods section, we assume a locus with two alleles, A and T (with allele frequencies P_A_ and P_T_, respectively), yet this of course applies to all possible nucleotides.

**Figure 5.**
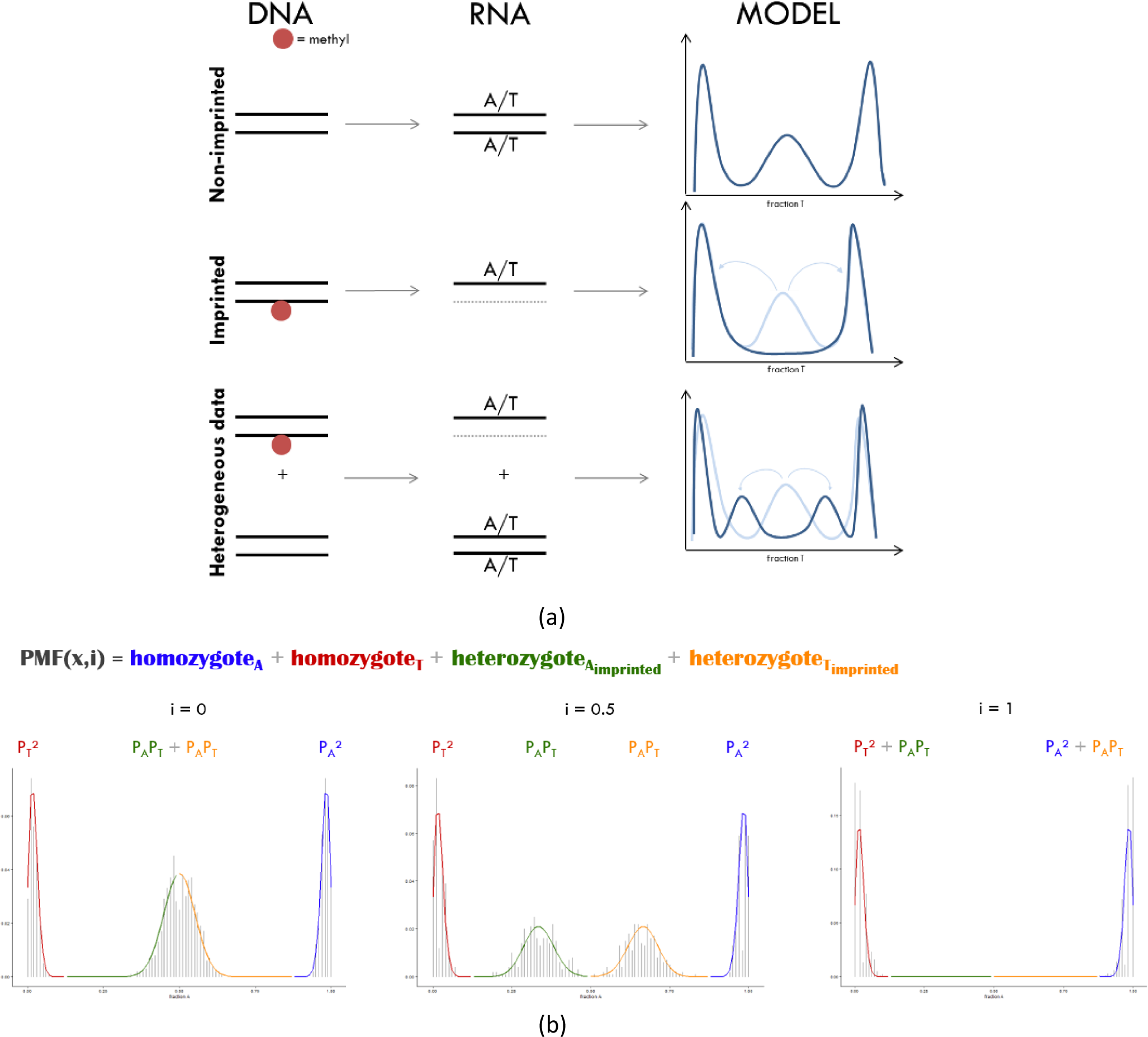
Graphical representation of rationale of the PMF. (a) The PMF is defined as a mixture model of genotype-dependent binomial distributions and describes the probability of observing specific RNA-seq coverages for each allele for a specific SNP locus. In these binomial probabilities, sequencing error rates, degree of imprinting (i) as well as the specific genotype are taken into account. For non-imprinted loci, the PMF results in two homozygous peaks and one heterozygous peak. For imprinting, on the other hand, no heterozygous can be detected on RNA-level and this peak is hence eliminated. Heterogeneous data leads to the detection of partial imprinting. (b) PMF for different degrees of imprinting. In this mixture model, the genotype-dependent binomial distributions have weights corresponding to their Hardy-Weinberg theorem derived expected chances.

Note that the same is true for enrichment-based sequencing data. Indeed, monoallelic histone modifications as well as monoallelic DNA methylation lead to ChIPseq and MethylCap-seq data, respectively, that is no longer in HWE, e.g.^27^.

To enable screening for loci featuring imprinting, a probability mass function (PMF) describing the probability of observing specific coverages for each allele for a specific SNP locus was developed. As the probabilities depend on the underlying genotypes, the PMF was created as a mixture model of genotype-dependent binomial distributions with weights corresponding to the probabilities under Hardy-Weinberg (Figure 5(b))^52^. Sequencing error rates (median per chromosome) are here taken into account. Subsequently, maximum likelihood estimation was used to estimate the degree of imprinting (i) and a likelihood ratio test was constructed to detect significant imprinting. A detailed discussion of the different elements of this PMF and the analytical framework can be found in Supplementary Methods, Section 2. All analyses were performed in R (v3.3.2)^53^, scripts are available upon request.

### 5. Detection of differential imprinting

Next to the detection of imprinted loci, we also examined possible deregulation of imprinting in cancer. This was done by testing for re-expression of the silenced allele, here termed differential imprinting. When occurring in non-normal samples (here cancer), this is coined loss of imprinting (LOI). Briefly, ratios of the lowest allele count over the highest allele count (i.e. R_i,s_ for SNP *i* and samples) are calculated for each single SNP, over all samples. These ratios are sorted per SNP in an ascending order separately for control and tumour samples. As the lowest ratios are expected for homozygous samples (ratios theoretically equal to 0, yet slightly higher due to the presence of sequencing errors), one can consider samples with the highest 2P_A_P_T_ ratios as putative heterozygous samples for that specific locus. In practice: samples with rank higher than round(sample_size*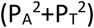) are considered as the heterozygous samples for a specific locus. After determining the mean ratio of these heterozygotes (Ri,stumour and Ri,control), parameter Ri,dw is calculated as the difference between these values, i.e. R_i,diff_ = R_i,stumour_ - R_i,s_ control. Upon random assignment of the tumour and control labels to the present samples by permutation, 10,000 random values of R_i,diff_ are simulated to generate a null distribution. Loci with an FDR-adjusted p-value smaller than the nominal 10% FDR level were concluded to be differentially imprinted between control and tumour samples (Supplementary Methods, Section 2.f). We were solely interested in LOI in cancer and hence we exclusively tested for higher ratios in tumour compared to control data. The different breast cancer subtypes were also tested for LOI (though the Normal-like subtype was not studied here, as only 8 samples were available) in which the p-value was corrected over all samples. Here, it should be noted that considering the ratios allows for detecting differences independent of alterations in expression levels of imprinted genes, which are also prominent in breast cancer^54^.

Subsequently, for loci featured by LOI, the latter was linked to survival with two different strategies. Firstly, the same allelic ratio as before was defined. Samples were once called LOI if the R_i,s_ was higher than 0.2 and once if it was higher than 0.5. Only SNPs with at least 5 LOI samples and samples for which the alternative allele was detected at least twice were retained. A Cox proportional hazards model was used to link survival to this categorical LOI variable, while adjusting for age. Secondly, a continuous LOI variable, which was defined as the allelic ratio, was also associated with survival, again adjusting for age with a Cox proportional hazard model. To anticipate assumption violations, a null distribution was constructed by 10,000 permutations by randomly shuffling the ratios over the samples. Loci with an FDR-adjusted p-value ≤ 0.1 were called significant. The analyses were also performed on solely the putative heterozygous samples, meaning that the 2P_A_P_T_ fraction of samples with the highest allelic ratios were used (see higher).

Differential imprinting of the alternative allele (lowest expressed allele) was analysed as well to assess whether expression of the normally silenced allele was higher in tumour data. Here, counts per million (CPM) reads of the alternative allele count were calculated as described in the next paragraph. The same permutation test yet based on the logCPM-values rather than ratios, was performed on the SNPs showing significant LOI.

### 6. Detection of differential expression

DE analysis was performed to further evaluate deregulation of imprinted genes in cancer. EdgeR normalisation factors were calculated from the breast cancer RNA-seq expression count file downloaded from firebrowse.org^55^. Afterwards, CPM-values for the imprinted SNPs were computed with these normalisation factors and library sizes (available for 100 control samples and 469 tumour samples: 87 BL, 54 HER2, 210 LumA, 111 LumB and 7 NL). EdgeR-based DE analysis was developed to increase power, at the cost of several assumptions. As the sample size and thus power is sufficiently high in the case at hand, we opted to use more robust standard non-parametric methods. Differential expression in control versus tumour samples was analysed with a Wilcoxon Rank Sum test for detected SNPs as well as the corresponding genes (sum of the CPM-values of the matching SNPs were used). To test for DE in the different breast cancer subtypes, a Kruskal-Wallis test and Dunn’s post-hoc test were performed on the CPM-values of the varying subtypes.

### 7. Analysis of HM13

Infinium HumanMethylation450k data and exonic expression data of HM13 were downloaded from firebrowse.org^55^. DE was analysed for 14 exons and 10 methylation probes (listed in Joshi *et al.*, with additionally cg18471488 as identified using MEXPRESS in the breast cancer population^28,29^) located in HM13. logCPM-values were calculated for the exonic data, normalised as described in the previous section. RPKM values, available from firebrowse as well, were used for additional verification and consistently yielded the same conclusions. Comparison between normal and tumour samples was done with a Wilcoxon Rank Sum test, while a Kruskal-Wallis test and Dunn’s post-hoc test were used for the breast cancer subtypes. Exon data were available for 468 tumour samples (85 BL, 54 HER2, 211 LumA, 111 LumA and 7 NL) and 100 control samples, whereas 450k Infinium methylation data could be retrieved for only 84 control and 207 tumour samples (35 BL, 14 HER2, 107 LumA, 46 LumB and 5 NL).

Subsequently, methylation and expression data were correlated to identify which methylated locus might control expression. Expression as well as methylation data were only available for 72 control and 192 tumour samples. A Spearman correlation test was performed for detecting correlation between the β-values of the 10 probes and i) logCPM-values of the whole HM13 gene (gene counts obtained from firebrowse.org^55^ and normalised with EdgeR) and ii) logCPM-values of the 3^rd^ exon of transcript 4 (as most of the imprinted SNPs were located in the neighbourhood of this exon). Again, RPKM values were successfully used for verification.

### 8. Quality control

To additionally verify the imprinting and deregulation results, corresponding WES data (BAM files) were downloaded from TCGA to obtain the underlying genotypes. However, only for 93 control samples and 464 tumour samples WES data were available. Concordance between WES and RNA genotypes was examined to validate the quality of genotyping with RNA-seq data (Supplementary Results, Section 3.c).

## AUTHOR CONTRIBUTIONS

TG and SS contributed equally to this work. The statistical approach was conceived by TDM and developed by TDM, TG and SS with biological and technical insight from WVC and OT; general strategy:

TG, SS and TDM, data pre-processing & management: TG, SS and JG. Implementation and data analyses were done by TG and CAV under supervision of SS and TDM. Additional validation was done by SS and CAV with contributions from TG and TDM. TG and SS wrote the manuscript with input from JG, CAV, TDM, WVC and OT. All authors read and approved the final text.

